# 100,000 years of gene flow between Neandertals and Denisovans in the Altai mountains

**DOI:** 10.1101/2020.03.13.990523

**Authors:** Benjamin M Peter

## Abstract

The Siberian Altai mountains have been intermittently occupied by both Neandertals and Denisovans, two extinct hominin groups^1,2^. While they diverged at least 390,000 years ago^3,4^, later contacts lead to gene flow from Neandertals into Denisovans^5,6^. Using a new population genetic method that is capable of inferring signatures of admixture from highly degraded genetic data, I show that this gene flow was much more widespread than previously thought. While the two earliest Denisovans both have substantial and recent Neandertal ancestry, I find signatures of admixture in all archaic genomes from the Altai, demonstrating that gene flow also occurred from Denisovans into Neandertals. This suggests that a contact zone between Neandertals and Denisovan populations persisted in the Altai region throughout much of the Middle Paleolithic. In contrast, Western Eurasian Neandertals have little to no Denisovan ancestry. As I find no evidence of natural selection against gene flow, this suggests that neutral demographic processes and geographic isolation were likely major drivers of human differentiation.

## Main text

The discovery of Denisovans is one of the early successes of the burgeoning field of ancient DNA^3,6–8^. Denisovan remains have been retrieved from Denisova Cave (Siberia, Russia)^3,6–9^ and a putative specimen has been reported from Baishiya Karst Cave (Xianhe, China)^10^. Most insights into Denisovans are based on the sole high-coverage genome of *Denisova 3^3^*: She was closely related to the Denisovans that interacted with the ancestors of present-day East Asians, but only distantly related to another population that interacted with the ancestors of present-day South-East Asians^5,7^. Much less is known for the three other Denisovans for which low-coverage genetic data has been retrieved (*Denisova 2, Denisova 4* and *Denisova 8*)^8,9^, where substantial contamination by present-day human DNA precluded detailed nuclear genetic analyses **(Figure 1a)**. Mitochondrial analyses revealed that *Denisova 4* differs at just two positions from *Denisova 3*, in contrast to the much more diverged lineage in the earlier *Denisova 2* and *Denisova 8* genomes^8,9^.

**Figure 1:**
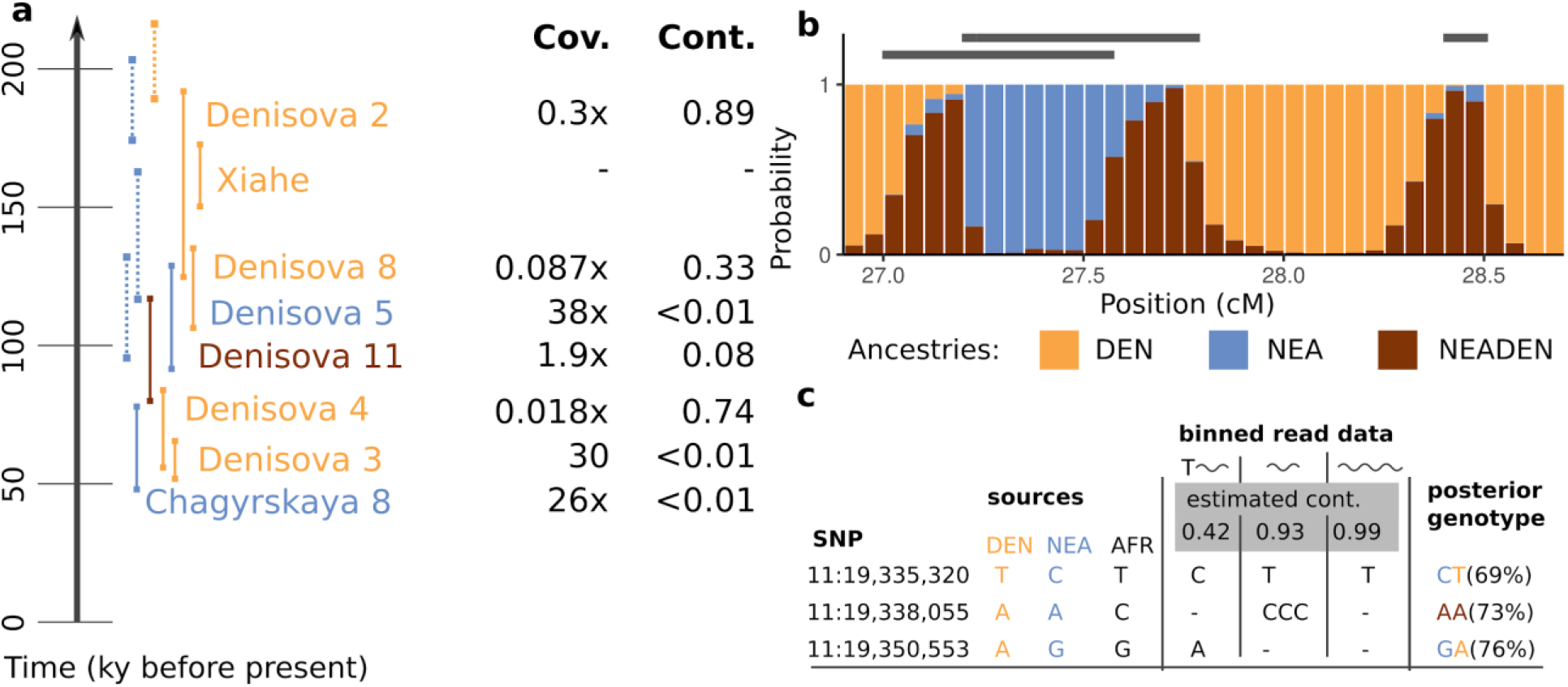
**a:** Archaic genetic data from the Altai mountains. Solid lines give confidence intervals for dates of specimens^1,2,13^, dotted lines layer ages for DNA retrieved from Denisova cave sediments^2^. Xiahe is the only Denisovan not from the Altai and is added as a reference. For each sample, average genomic coverage and modern human contamination estimates are displayed. **b**: Schematic of local ancestry model used in admixfrog. Shown is a 2cM region of a simulated Denisovan chromosome with three introgression fragments (grey bars). The barplot depicts the posterior decoding obtained using admixfrog from low-coverage (0.1x) data. Heterozygous ancestry is called in regions where only one introgression fragment is present; homozygous Denisovan ancestry is called where they overlap. **c**: Overview of the genotype likelihood model, based on three SNPs in a heterozygous region of *Denisova 2*. We display the allele in two source populations (Denisovans and Neandertal) as well as Sub-Saharan Africans (AFR) as a proxy for the contamination source. Read data is split into three bins based on whether sequences carry a deamination (*T*~) and sequence length (~ vs ~~). Letters give the number of sequences with a particular base overlapping this position. The resulting posterior genotype shows that read bins with high contamination rates are efficiently downweighted, resulting in a posterior reflecting the archaic ancestry.

The Altai region has also been occupied by Neandertals, as evidenced by hominin remains, artifacts and DNA from multiple sites^11–13^. This co-occupation history resulted in gene flow from Neandertals into Denisovans: Comparisons of the high-coverage *Denisova 5* (“Altai”) Neandertal^5^ with *Denisova 3* revealed a small proportion (0.5%) of net gene flow from Neandertals into Denisovans^5^. Direct evidence of contact was provided by the discovery of *Denisova 11*, the offspring of a Neandertal mother and a Denisovan father^6^. Additionally, tracts of homozygous Neandertal ancestry in this genome suggest that the father had additional Neandertal ancestors several hundreds of generations ago.

While early methods to detect gene flow from ancient DNA used genome-wide summary statistics^5,14^, inference may also be based on directly detecting genomic regions where an individual harbors ancestry from a different population. Approaches using these “admixture tracts” are more sensitive when overall levels of gene flow are very low^5^, and have provided much evidence about when and where gene flow between archaic and modern humans happened^15,16^, and about the functional and phenotypical impact of that gene flow^17^. However, most current methods to infer admixture tracts assume high-quality genotypes^15,18^ and are thus not applicable to the majority of ancient genomic data sets, which are frequently low-coverage, and contaminated with present-day human DNA^19,20^.

### The Admixfrog Model

As recently introgressed tracts can stretch over thousands of informative SNPs, combining information between markers allows inference from low-coverage genomes^21–23^. Here, I combine a Hidden Markov Model for local ancestry inference with an explicit model of present-day human contamination in a program called admixfrog **(Methods, Supplement 1).** Briefly, I assume that the analyzed *target* individual has ancestry from two or more *sources*, that represent potentially admixing populations. The sources are represented by high-quality genomes; in all applications I use two high-coverage Neandertals (NEA)^4,5^ and the high-coverage *Denisova 3* (DEN)^3^ genomes. Admixfrog infers the tracts of the target individual’s genome that originated from each source **(Figure 1b)**. In contrast to most previous approaches, I use a flexible empirical Bayes model to estimate all parameters directly from the data, thus alleviating the dependence on simulations or strong modelling assumptions about admixture times or past population sizes, which may introduce unwanted biases^24,25^. This local ancestry model is combined with a genotype likelihood model that incorporates present-day human contamination **(Figure 1c)**, taking into account that contamination rates are influenced by technical covariates such as sequence lengths^26^, terminal deaminations^27^ or differences between libraries.^28,29^

### Validation

I validate admixfrog using simulations on scenarios of gene flow from Neandertals into Denisovans and modern humans **(Methods, Extended Data Figs. 1–3).** In cases without admixture, tracts longer than 0.1cM are inferred with precision of 96% even for 0.03x genomes, relatively independent of sample age. Contamination decreases the performance, but particularly in scenarios of gene flow between archaics, fragments longer than 0.2cM are highly accurately inferred. I also use experiments modifying real data^5,8,30^ to evaluate admixfrog under more realistic conditions, to compare it to other methods, and to assert its robustness to parameter choices (recombination map, SNP ascertainment, sources, etc.), **(Extended Data Fig. 4, Methods)**^8^. In most tested cases, I find that admixfrog produces comparable results to those obtained from high-coverage data, and that long introgression tracts can be recovered even in ultra-low coverage genomes. This suggests that the program is well-suited for the analysis of ancient genetic data from both present-day and archaic hominins.

### Recent Neandertal ancestry in early Denisovans

The genomes of the two oldest Denisovans, *Denisova 2* and *Denisova 8^8,9^* are both highly contaminated (**Figure 1a, Extendend Data Fig. 5ab**), with estimated coverage by endogenous molecules of 0.030x and 0.087x, respectively. As even sequences starting with deaminations^31^ have significant amounts of contamination **(Extended Data Fig. S5ab)**^8,9^, previously used filtering techniques would fail^26^. Despite this, admixfrog identifies 212.6 cM (173Mb) of Neandertal ancestry in *Denisova 2,* and 258 cM (210Mb) in *Denisova 8* **(Figure 2, Extended Data Table 1).**

**Figure 2:**
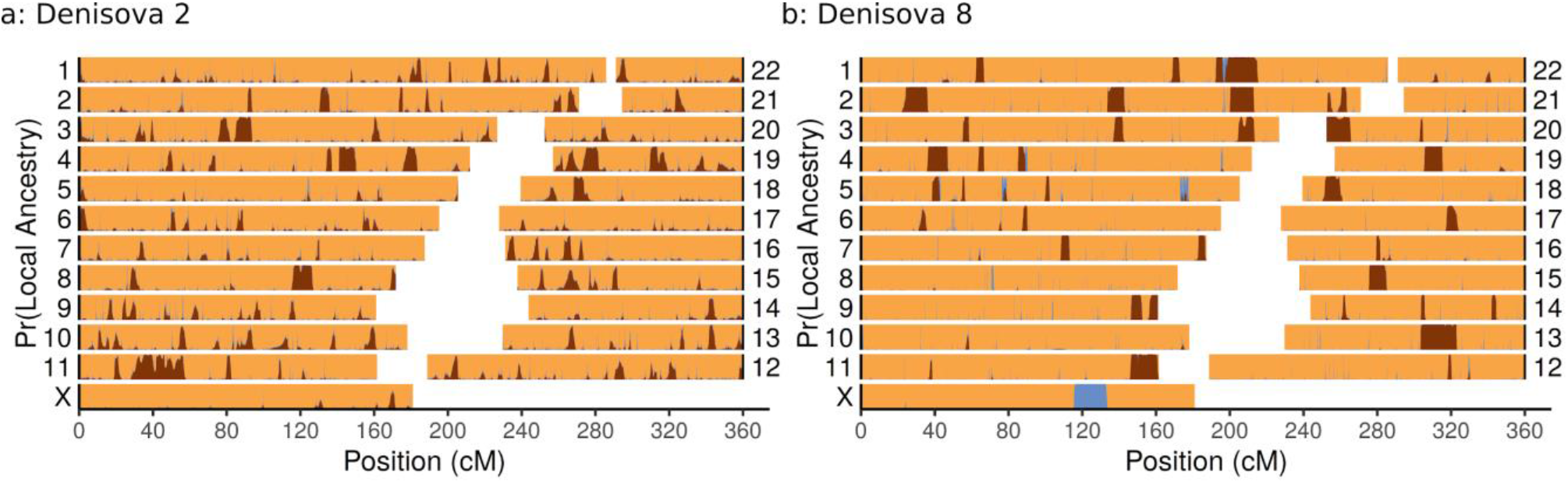
Neandertal ancestry in Early Denisovans. We show the admixfrog posterior decoding of **a**: *Denisova 2* and **b**: *Denisova 8*. Homozygous Denisovan ancestry, homozygous Neandertal ancestry and heterozygous ancestry are in orange, blue and brown, respectively.

The longest inferred tract for *Denisova 2* is located at chr11:18,791,748-36,393,966 (hg19), and has a recombination length of 25.7 cM. To confirm this finding, I perform a validation analysis insensitive to modern human contamination **(Extended Data Fig. 7a):** The data is restricted to SNPs where *Denisova 2* reads carry an allele never found in modern humans, and where either Denisovans or Neandertals, but not both match the non-human allele seen in *Denisova 2*. At 45 of these 81 sites, *Denisova 2* carries the Neandertal allele, which is consistent with the 50% expected in a region of heterozygous Neandertal-Denisovan ancestry. The average length of Neandertal ancestry tracts in *Denisova 2* suggests that most Neandertal ancestry dates to around 1,500 years prior to when *Denisova 2* lived (50±10 generations, mean ± 2sd, generation time of 29 years, **Extended Data Table S1**), but the longest tract is likely younger (14.1±14 generations), hinting at more recent Neandertal ancestors. Results for *Denisova 8*, are qualitatively similar, but the higher coverage of 0.087x allows more accurate estimation of fragment boundaries **(Figure 2b).** Overall, *Denisova 8*’s Neandertal ancestry is more recent (22±6 generations), as evidenced by a 23.7Mb (22.5cM) tract on chr1:179,807,948-203,527,526, and seven other tracts longer than 10cM, including one on the haploid X chromosome (chrX:114,752,520-124,257,661, **Extended Data Fig. 7b**). The similar amount and tract lengths of Neandertal ancestry in *Denisova 2* and *Denisova 8* raise the possibility that they resulted from the same gene flow event, in particular since the stratigraphic location of *Denisova 2* cannot be established conclusively, and so its age might be close to *Denisova 8^1,2^*. To test this hypothesis, we compare the locations of Neandertal ancestry tracts between the genomes. If the tracts in both specimens traced back to the same introgression event, their spatial location should be correlated^32^. However, this is not the case (Fisher’s exact test, p=0.56), suggesting that they belonged to different populations with distinct Neandertal introgression events. The finding that the locations of introgressed tracts are uncorrelated also rules out the potential issue that gene flow into the reference *Denisova 3* might be confounded with gene flow into the earlier Denisovans, as such a bias should be present in both genomes and thus cause a correlation between introgression tract locations. Such a signal is indeed observed in the HLA region on chromosome 6 **(Extended Data Fig. 5)**, which is unsurprising given the age of haplotypes there. Similarly, the tract locations are also not significantly correlated with the homozygous Neandertal-ancestry tracts of *Denisova 11,* if the HLA region is removed **(Extended Data Fig 8a)**, (Fisher’s exact test; p=0.05 and p=0.09 for *Denisova 2* and *8*, respectively).

### Gene flow into Neandertals

In addition to the gene flow from Neandertals into Denisovans described here and previously, we also identify recent Denisovan ancestry in two Neandertals^5,33^ from the Altai Mountains. Although the overall proportion of inferred Denisovan ancestry in *Denisova 5* is small (0.15%), six out of the 15 identified tracts exceed 1 cM in length **(Figure 3a, Extended Data Fig. 7c).** The longest fragment is a 2.0Mb (2.18cM) fragment on the X-chromosome (chrX:136,505,565-138,501,953). The length of these fragments suggests that gene flow happened 4,500±2,100 years before *Denisova 5* lived. A lower total of 3.8 cM (4.8Mb) of Denisovan introgressed material is found in *Chagyrskaya 8*, a more recent Neandertal from the Altai mountains^33^. The inferred tracts are small, with the longest tract measuring 0.83 cM **(Extended Data Fig. 7d)**, suggesting that this gene flow happened several tens of thousands of years before *Chagyrskaya 8* lived. In contrast, little to no Denisovan ancestry is detected in eight Western Eurasian Neandertal genomes^4,20,29^ dating from between 40,000 and 120,000 years ago **(Extended Data Fig. 8, Extended Data Table 1, Supplementary Table 1)**. In three of these genomes (*Goyet Q56-1, Spy 1 and Les Cottes*), the centromere of chromosome 10, a region implicated in gene flow between archaic and modern humans^34^, is identified as introgressed from Denisovans. Thus, while Denisovan alleles survived for many millennia in Altai Neandertal populations, little to none of that ancestry made it into later Neandertal populations in Europe.

**Figure 3:**
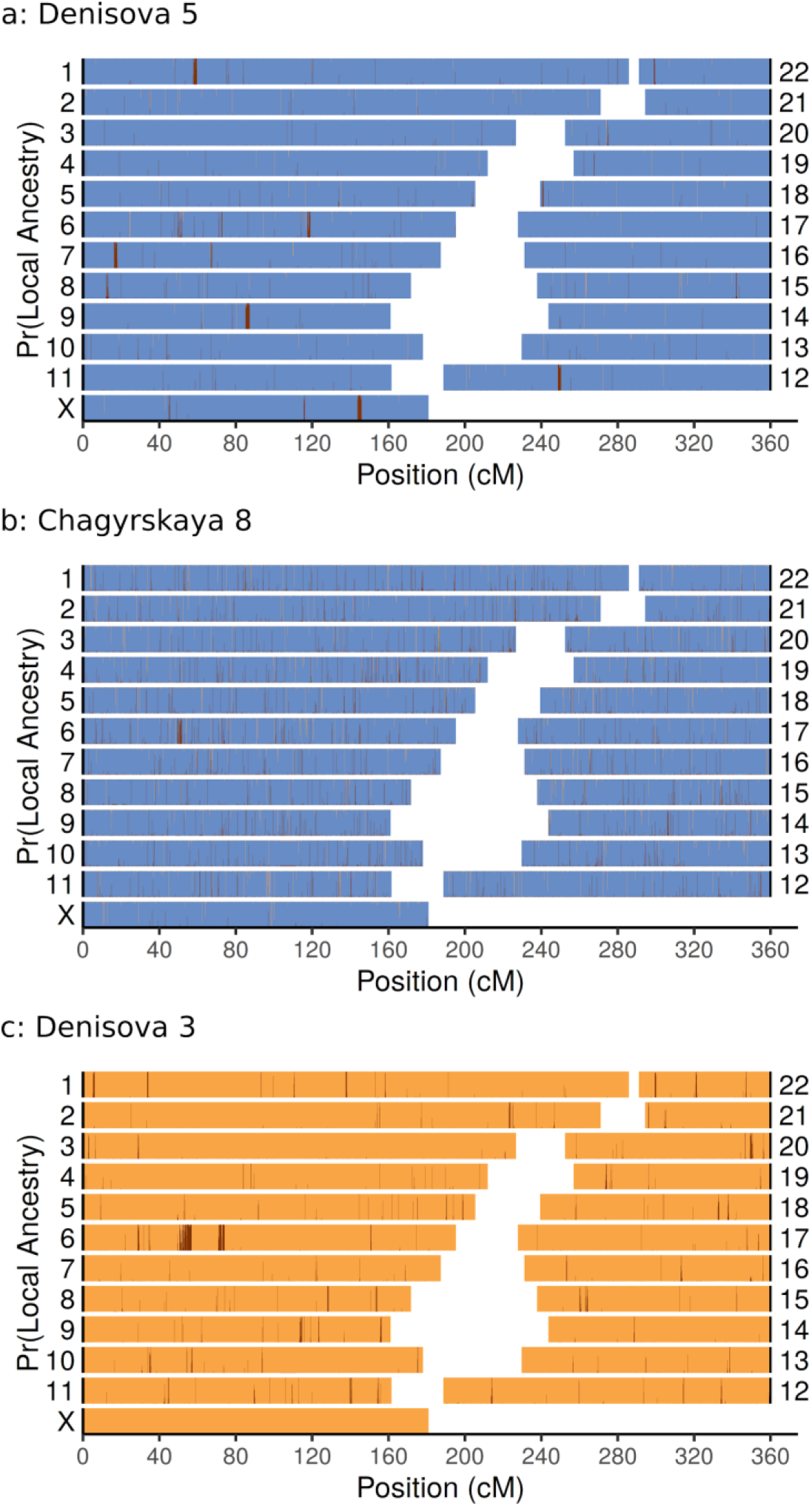
Evidence for later admixture. We show the admixfrog posterior decoding of a: *Denisova 5*, b: *Chagyrskaya 8* and c: *Denisova 3* (using fixed priors). Homozygous Denisovan ancestry, homozygous Neandertal ancestry and heterozygous ancestry are in orange, blue and brown, respectively.

### Gene flow into late Denisovans

As *Denisova 3* is the sole reference for Denisovan ancestry, and coverage for the other late Neandertal, *Denisova 4,* is too low (<0.005x, **Extended Data Figs. 6f, 8b)**, I screen for Neandertal ancestry in *Denisova 3* using a modified analysis using a fixed prior **(Methods).** This analysis amounts to scanning for large genomic regions where Denisova 3 has a large number of heterozygous sites, but few homozygous differences to Neandertals. We validate this analysis using two other high-coverage Neandertals **(Extended Data Fig 9, Supplementary Table 1)**, finding that results for these genomes are more noisy than the standard analysis, but qualitatively similar. to the standard analysis. This procedure discovers a total of 58 Neandertal introgressed fragments in *Denisova 3*, amounting to 21.5 Mb (0.4% of the genome), roughly double the amount found in *Denisova 5* **(Figure 3c**, **Extended Data Fig. 9g)**.

### Diversification despite gene flow

The presence of ancestry tracts in all analyzed genomes from the Altai suggests that gene flow between Neandertals and Denisovans was prevalent, and occurred recurrently over up to 100,000 years. For the first time, we demonstrate that Altai Neandertals have Denisovan ancestry, showing that their offspring must have been able to produce fertile offspring with both populations. As there is also no evidence for reduced introgressed ancestry on the X-chromosome (p=0.47, permutation test), no association of introgressed regions with levels of background selection^35^ (p>0.14, permutation test) and no evidence that introgressed tracts correlate with any functional annotation category (GO-enrichment analysis^36^, hypergeometric test, p>0.05 for all categories), and no significant association of introgression tracts with regions depleted for Neandertal ancestry in modern humans (p=0.317, permutation test), there is no evidence of negative fitness consequences of Neandertal-Denisova matings.

This suggests that the genetic and morphological^10,37^ differentiation between Neandertals and Denisovans is substantially due to neutral processes, i.e. geography. A plausible scenario is one where the Altai Mountains are part of a relatively stable hybrid zone, that persisted through multiple warmer and colder periods^2^. While matings might have been locally common, migrations from the Altai to Europe were likely scarce, as evidenced by the almost complete absence of Denisovan ancestry in European Neandertals. A similar scenario seems likely for Denisovans; as the later *Denisova 3* has much less Neandertal ancestry than the earlier *Denisova 2* and *Denisova 8*, it must have received substantial ancestry from a reservoir Denisovan population with little Neandertal ancestry. Similarly, the finding that the location of introgression tracts are independent between genomes suggests that the Denisovan occupation in the Altai region was not continuous. More speculatively, findings of early gene flow between modern humans and Neandertals^38,39^ perhaps suggests that a similar relationship of occasional gene flow followed by local extinctions also existed between modern humans and Neandertals, before early modern humans migrated out of Africa and displaced the resident Eurasian hominins.

## Methods

### The admixfrog algorithm

Inference is based on version 0.5.6 of the program admixfrog, which is available from https://github.com/BenjaminPeter/admixfrog/. Full details and derivation of the algorithm are given in **Supplemental Text 1**. Briefly, a *target* individual is modelled as a mixture of two or more *sources*, informed by the allele frequencies in a sample of high-quality genomes. The model is implemented as a Hidden Markov Model, where the hidden states are all diploid combinations of hetero- and homozygous states from the sources. Compared to similar approaches^22,40^, admixfrog mainly differs in that i) almost all parameters are directly learned from the data, ii) contamination and uncertainty due to low-coverage is modelled explicitly using a genotype-likelihood model and iii) multiple ancestries can be distinguished.

Admixfrog models genetic drift and sampling uncertainty using a modified Balding-Nichols model^41^, and thus does not require phased genomes as input. For each source, two nuisance parameters, τ and *F,* measure genetic drift before and after admixture. Two additional parameters, *a*_0_ and *d*_0_ are Beta-distribution priors and reflect how well the available sample from the source reflects the population allele frequency. This local ancestry model is combined with a genotype likelihood model that incorporates contamination. As contamination rates are expected to differ based on covariates such as the presence of terminal deaminations^27^, read lengths^26^, or library^28,29^. Reads are grouped into discrete bins based on these covariates. Contamination rates are then independently estimated for each bins.

### Data processing and references

Admixfrog requires a set of high-quality reference panels consisting of one or multiple high-quality genomes, and ascertainment to a pre-specified set of single nucleotide polymorphisms (SNPs) that are variable between these references. These references are then either used as *sources,* i.e. potential donors of admixed material, or as putative contaminants. In all analyses presented here, the following references are used: *AFR*, consisting of the 44 Sub-Saharan Africans from Simons’ Genome Diversity Panel (SGDP)^42^, as a proxy for modern humans and contaminants. To model Denisovan ancestry, I use the high-coverage Denisova 3^3^ genome *(DEN)*, and for *NEA,* reflecting Neandertal ancestry, I use the two high-coverage Vindija 33.19 and Denisova 5 (“Altai”) Neandertals^4,5^. I also use the chimpanzee (panTro4) reference genome allele as a putative archaic allele. For supplementary analyses, I also use *Vindija 33.19* (*VIN*), *Chagyrskaya 8* and *Denisova 5* (*ALT*) genomes individually, and use SGDP Europeans *(EUR)*, SGDP East Asians (EAS) or 1000 genomes Africans *(AFK)* for modern human ancestry. Throughout, I use a bin size of 0.005 cM for local ancestry inference, based on a linearly interpolated recombination map inferred from recombination patterns in African Americans^43^, obtained from https://www.well.ox.ac.uk/~anjali/AAmap/.

For the *target* samples I start with aligned reads stored in bam files, obtained from the authors of the respective publications^3–6,8,9,33^. In all cases, I filtered for reads of lengths of at least 35 base pairs; and mapping quality >= 25, variable positions matching a C->T substitutions in the first three positions of the read, or a G->A substitutions at the end of a read were discarded. Only 2,974,930 positions known to be variable between the three high-coverage Neandertals and Denisova 3 are considered. Sites were pruned to be at least 50bp (415,546 SNPs removed), and 0.0001 cM (261,587 SNPs removed) apart, resulting in 2,297,797 SNPs used for analyses.

The exact command run is

~~~
admixfrog --infile {infile} --ref ref_hcneaden.csv.xz -o {outname} --states NEA_DEN --cont-id AFR --
ll-tol 0.01 --bin-size 5000 --est-F --est-tau --freq-F 3 --freq-contamination 3 --e0 0.01 --est-
error --ancestral PAN --run-penalty 0.2 --max-iter 250 --n-post-replicates 200 --filter-pos 50 --
filter-map 0.000
~~~

Where infile, and outname are substituted by the respective files for each analyzed specimen. The pipeline used to create all data except the simulations is available at https://github.com/BenjaminPeter/admixfrog/tree/master/pipeline.

### Dating information

No new dating is performed in this study, but.dating information is important for context. For most specimens, dates are taken from the papers describing the genetic data. For Denisova Cave specimens, I use Bayesian dates^1^, and for sediments the layer dating results^2^. For Vindija 33.19 and Chagyrskaya 8, I report results from the most recent reports^13,44^.

### Covariates for contamination rate estimation

Admixfrog co-estimates contamination from an assumed contaminant population, taking into account that contamination rates may vary with covariates such as read length^26^, terminal deaminations^27^ and library preparation methods^28^. For this reason, I group reads into discrete sets according to i) the library they are coming from, ii) whether the read has a terminal deamination or not and iii) the length (in bins of 15 bp, i.e. bin 1 contains reads with lengths between 35 and 49bp, bin 2 between 50 and 64, etc). Contamination and sequencing error rates are then estimated for each of those sets of reads, independently.

### Calling fragments

By default, admixfrog returns a posterior decoding, i.e. the probability that a given bit of the genome has a particular local ancestry. To call tracts, I combine adjacent fragments using a simplified Needleman-Wunsch algorithm^45^, using the scoring function *S*_*i*_ = max[*S*_*i-1*_ + log(*p*_*i*_ + *r*), 0]; *S*_0_ = 0, where *p*_*i*_ is the posterior probability of being in any target state. In particular, I focus most of our evaluation on admixfrog’s ability to call introgressed ancestries, regardless of whether it is present as homozygous or heterozygous. Then, I can simply sum the posterior probabilities for the states NEA and NEADEN at each position. The parameter *r* models how generous I are at bridging short gaps; a high value of *r* tends to merge adjacent fragments, whereas a low value of *r* might split up fragments. All analyses use *r=0.4*.

Based on *S*_*i*_ fragments are called using a simple backtracking algorithm:

while any *S*_*i*_ > 0:

- Find end position: *i*_end_ for which *S*_*i*_ is maximized
- Find start position: *i*_start_ max_*i*_ *i* = 0, *i* < *i*_max_
- Set p[*i*_start_ : *i*_end_] to 0
- Recalculate *S*_*i*_ = max[*S*_*i-1*_ + log(*p*_*i*_ + *r*), 0]

### Estimating admixture time

Under a simple model of archaic admixture at a single time point, the lengths of introgression tracts follow an exponential distribution with rate proportional to the admixture time. The maximum likelihood estimator for the mean admixture time (in generations) given *n* introgressed fragments *L*_i_ at least *c cM* long, is, 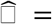 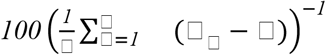. Throughout, I use a cutoff of *c*=0.2 cM, as our simulations indicate that the vast majority of fragments are detected above this length for recent admixture **(Extended Data Figures S1, S2)**.

To show that the most recent Neandertal ancestor of *Denisova 2*, was likely recent, I use a simulation based estimator. I simulate a genome of size *G*=3740 cM. after *t* generations, it will have recombined *tG* times, and a proportion *p*=0.03 of fragments will be of Neandertal ancestry. I simulate 10,000 genomes each at each time point from one to 100 generations ago, and record the longest introgressed fragment *L*. For all fragments of a given lengths *L*, I record the mean, 5% and 95% quantile.

### Modified analysis using fixed site frequency spectrum prior

As Denisova 3 is the only high-quality Denisovan genome available, the standard algorithm overfits the Denisovan ancestry of *Denisova 3*. To avoid this, I flatten the allele frequency priors by setting the site-frequency-spectrum priors *a*_0_ = *d*_0_ = 0.1 instead of estimating these parameters from the data. This has the effect that the DEN source becomes uniformly less similar to *Denisova 3,* and allows the identification of regions that are much more similar to Neandertals than the genomic background. I validate this approach using the the high-coverage *Denisova 5* and *Vindija 33.19* genomes, using only a single Neandertal and Denisovan genome each as references.

### Functional annotation and selection analysis

To investigate the functional consequences of gene flow between Neandertals and Denisovans, I perform a number of tests where I compare whether introgression tracts are significantly associated with a number of genomic features. Null distributions are obtained by randomly shuffling the observed fragment location 1,000 times.

#### B-statistics^35^

B-statistics are a measure of local background selection, and have been shown to be positively correlated with Neandertal ancestry in modern humans^25^. I perform a similar analysis by annotating each bin used for analysis with its mean B-statistic, lifted over to hg19 coordinates. I then calculate the proportion of introgressed material from all analyzed genomes in five quantiles of B-statistics.

#### Overlap with deserts of Neandertal introgression

I compare introgression tracts with regions where modern humans are deficient of Neandertal ancestry (‘deserts’)^46^. Four of these deserts have evidence of Neandertal ancestry in Denisovans, with Denisova 2, Denisova 3 and Denisova 11 having one, and Denisova 8 having two fragments overlapping deserts. For two of the deserts (the ones on chromosomes 3 and 18) I do not find any overlapping introgressed fragments. Overall, there is no evidence that Neandertal introgression is more or less frequent than expected by chance (p=0.32).

#### X-chromosome

When resampling all introgressed tracts, I assert the proportion of introgressed material on the X-chromosome. I do not find an enrichment, but note that the confidence intervals are very wide (0.035-0,223);

#### Functional enrichment

I perform functional enrichment using a hypergeometric test as implemented in the GOfuncR package ^36^. Enrichment is performed by i) inferring all genes contained in an introgressed region in any individual and ii) performing functional enrichment against all GO-categories. After controlling for family-wise error rates, all categories are non-significant.

### Empirical Tests

To evaluate the performance of admixfrog under realistic conditions, I use computational experiments using two high-coverage and one low-coverage ancient genomes, under the premise that fragments should be reliably inferred using all available data. The genomes used are the ~45,000 year old Ust’-Ishim^30^, a modern human sequenced to 42x, the ~110,000 year old Denisova 5 Neandertal^5^ sequenced to 50x and the ~120,000 year old Denisova 8 genome. For ease of presentation, only one chromosome is plotted (chr1 for Ust-Ishim Denisova 8, chr9 for Altai), although the model fitting was done using the full genome. The basic strategy here is to perform a series of analyses where the default parameters mimic those used in the main data analysis. Each run then modifies one or multiple parameters to test its impact. For Ust’-Ishim, the analyses are modified by i) adding AFR as a proxy for modern human ancestry, and ii) ascertaining SNPs according to the archaic admixture array ^47^. The following scenarios are presented in **Figure S5**:

#### Previous methods

I compare our results with the approach of Fu et al.^30^, who simply plotted the location of SNPs where Africans are homozygous ancestral, and Denisova 5 carries at least one derived allele. I also compare the inferred fragments with those based on SNP-density in an ingroup, using a method proposed by Skov et al. ^48^, which uses a very different signal in the data but results in largely consistent calls **(Fig S5b)**.

#### Low coverage data

Lower coverage is achieved by downsampling the genomes by randomly discarding a fraction of the reads (using the --downsample option in admixfrog). In Denisova 5, 2%, 0.06%, 0.02% and 0.01% of reads are retained. For the Ust’Ishim genome, 100%, 1%, 0.25% and 0.025% of reads are retained **(Fig S5cd)**.

#### Contamination

I also performed analyses of the genome downsampled to 10% of the original coverage, adding between 5 and 80% contamination for all read groups, directly from the contamination panel **(Fig S5ef)**.

#### Parameter settings

For Denisova 5, I explore some different settings. In particular, I i) fit admixfrog using called genotypes^5^ rather than the genotype likelihood model (“GTs”), ii) I add two additional states for inbred Neandertal / Denisovan ancestry (“inbr.”), and I run analyses without estimating hyperparameters (“fixed”), and without an ancestral allele (“noanc”) **(Fig S5g)**.

#### Ascertainment schemes

As ancient data particularly from low-yield samples is frequently generated using capture enrichment^49^, I test a variety of SNP ascertainment schemes. Low-frequency or fixed SNPs have little impact on the likelihood, so ascertainments may be desirable to save memory even for shotgun data. I investigate four ascertainment schemes: i) the Archaic admixture array, containing 1.7M SNPs fixed between Africans and Denisova 5 / Denisova 3 (“AA”)^49^, ii) the 1240k array, which is widely used in the analysis of Neolithic and later human populations^49^, iii) the 3.7 M array, which is a combination of i and ii); iv) pANC, an ascertainment based on all segregating sites between Vindija, Altai Neandertal, Chagyrskaya 8 and Denisova 3 **(Fig S5h)**.

#### Sources

I also investigated the effect of different sources; either adding AFR as a source (“AFR”), or replacing the combined NEA ancestry with individual Neandertals (VIN/ CHA/ ALT) **(Fig S5ij)**.

#### Prior

The parameters *a*_0_ and *d*_0_ of the site-frequency-spectrum prior.

#### Bin Size

I also investigate the effect of changing the size of each bin from 0.005 cM to 0.002, 0.01 or 0.05 cM **(Fig S5kl)**.

#### Recombination map

Besides the African American map^43^ used for most analyses, I use physical distance (“none”), the deCode map (“deCODE”)^50^, and a hapmap map based on Yorubans (YRI)^51^ **(Fig S5op)**.

### Simulations

I use computer simulations to ascertain the accuracy of admixfrog under a number of scenarios. Simulations are performed in a coalescent framework using msprime 0.7.0^52^, which allows direct recording of which parts of the genome were introgressed. Throughout, I use a simple demographic model of archaic and modern humans, and replicate each simulation 20 times. The simulations are set up in a reproducible pipeline using snakemake^53^, available under https://www.github.com/benjaminpeter/admixfrog-sims

#### Simulation settings

Here, I outline the baseline model used for all simulations. I assume a model where hominins split 6 Million years ago (ya) from the primate ancestor. 600kya the early modern humans split from the common ancestor of Neandertals and Denisovans, which themselves split 400kya. Within the modern human clade, Africans split from Non-Africans 70kya, and Asians and Europeans split 45kya. Effective population sizes are set to 10,000 for Africans and 1,000 for archaic populations. For Non-Africans, the present-day population size is 10,000 but I assume a size of 2,000 from 70ky-10kya, to model the out-of-Africa bottleneck.

#### Gene flow

I model and investigate two gene flow events: Gene-flow from Neandertals into Europeans happened 50kya, and replaced 3% of genetic material. Gene flow between Denisovans and Neandertals is set to 120kya, and replaces 5% of genetic material. Both gene flows are assumed to occur within one generation.

#### Reference panel

Within this model, I create a reference panel that mimics the data used for the admixfrog analyses. In particular, I create a Neandertal source by sampling 3 Neandertals at 125kya, 90kya and 55kya, and a Denisovan source at 50kya, respectively. I sample further a reference panel of 20 diploid present-day Africans. A panel of 5 diploid European genomes is further simulated to model contamination. In addition, a single chromosome from the Chimpanzee is sampled as putative ancestral allele.

#### Data generation

For each scenario, I generate genetic data from *target* individuals, which are single diploid samples. Two diploid Denisovan samples each are taken at 110 kya 100 kya. 70 kya and 50kya, and two diploid early modern human samples each are taken 45kya, 30kya, 15kya and at the present time. I generate 20 chromosomes of size 50 MB each, assuming a constant recombination rate of 10^-8^. SNP are ascertained either to be variable in archaics (for scenarios looking at gene flow within archaics) or to be variable between archaics and modern humans (for scenarios investigating gene flow into modern humans). From the simulated target individual, read data is generated in one or multiple libraries independently, where each library *l* has a target coverage D*l* and a contamination rate *cl* For each polymorphic site *s* in each library, I assume that coverage is Poisson distributed: *C*_*sl*_ ~ Poisson(*D*_*l*_ [*c*_*l*_ *q*_*s*_ + *(1-c_l_)p_s_])*, where *p*_*s*_ and *q*_*s*_ denote the allele frequencies in the target individual and contaminant panel, respectively. In all tests including contamination, I simulated 10 libraries with different contamination rates between 0 and 90%, as indicated in **Extended Data Figures 1d, 2d**.

#### Running admixfrog in simulations

I run admixfrog using the following command (strings in curly brackets are replaced depending on the scenario), estimating all hyperparameters.

~~~
admixfrog --infile {input.sample} --ref {input.ref} -o {outfile} --states {state_str} --
cont-id AFR --ll-tol 0.001 --bin-size {bin_size} --est-F --est-tau --freq-F 3 --freq-
contamination 3 --e0 0.01 --est-error --ancestral PAN --max-iter 100 --n-post-replicates
100 --run-penalty 0.4
~~~

#### Evaluation of Simulations

The main purpose of admixfrog is the identification of genomic regions that are introgressed, and so I are interested under which condition I may expect to successfully recover these admixture tracts. As msprime simulations allow us to record when and which bits of an individual’s genome are introgressed from another population, I can compare the admixfrog results with the ground truth from the simulation.

In particular, I classify fragments as

1. **True positives** are fragments that are detect in some shape or form, I further subdivide them as
  a. **Strict true positives** are fragments that are correctly inferred as a single fragment
  b. **“Overlap”**-fragments occur when introgressed fragments on the two chromosomes of an individual overlap, so they cannot be distinguished under the admixfrog model
  c. **“Gap”**-fragments are introgressed fragments that are erroneously split into two or more different introgressed fragment
  d. **“Merged”** fragment are two adjacient introgressed fragments that are erroneously inferred as one
2. **False positives** are non-introgressed fragments that are inferred to be introgressed
3. **False negatives** are introgressed fragments that are undetected by admixfrog

Depending on the analysis, I are interested in the *precision,* (proportion of true positives among inferred fragments), which measures how much I can trust the detected fragments, the *sensitivity* (proportion of true positives over all true fragments), and the *proportion of true positives* (proportion of strict true positives + overlap over all true positives), which measures how often I split or merge fragments.

### Findings

I evaluate the performance of admixfrog in scenarios of admixture from Neandertals into Denisovans **(Extended Data Figures 1, 3)** and modern humans **(Extended Data Figures 2, 3)**.

#### Gene flow into modern humans

For modern humans, I assume a larger effective size of *N*=2,000, and admixture 50,000 years ago (generation time 25 years). Under such conditions, precision for 200kb tracts is above 95% in all scenarios, even at the lowest coverage of just 0.03x **(Extended Data Figure 1a)**. Sensitivity is more strongly impacted by coverage. While 2x coverage is sufficient for detection of admixture tracts in all scenarios, low-coverage fragments become harder to detect in older genomes. In the scenario where gene flow occurs just 5,000 years after admixture, around 25% of fragments of 0.5Mb length are missed at 0.03 coverage. At contamination levels below 20%, contamination levels are accurately estimated **(Extended Data Figure 1d),** and classification accuracy for fragments longer than 200kb remains very high. However, for higher contamination scenarios, the estimates become flattened, in that the differences between libraries are not correctly recovered, and more false positives are observed. As this is not the case when adding contamination from the “correct” contamination panel **(Extended Data Figure 4ef)**, this is likely due to genetic differences between the simulated contaminant (Europeans) and the one used for inference (Africans).

#### Gene flow into Denisovans

This scenario differs from modern humans in that the effective population size is smaller (*N*=1,000), admixture is older (120,000 years ago), and that the contaminant is more distinct from the sample (African contamination in a Denisovan individual). The smaller effective size results in high genetic drift (the two more recent samples are taken >2N generations after gene flow), and thus most fragments are short, and difficult to infer from low-coverage data, where sensitivity is close to zero **(Extended Data Figure 2)** and samples are frequently inferred as having no introgression at all. However, at higher coverages of 0.5x and 2x, introgressed fragments are detected and sensitivity approaches 50%. For the two sampling points closer to the admixture time, both precision and sensitivity are higher; and at coverages 0f 0.5x and 2x both sensitivity and precision are above 0.75 for fragments longer than 200kb. Thus, I find admixture between archaics is mainly detectable in the first few tens of thousands of years after gene flow, but I may struggle to detect older admixture. In contrast to the scenario where I simulate admixture into modern humans, I find that contamination rates are accurately estimated in all scenarios, but with a slight underestimate, most likely also due to the conservative misspecification of the admixing population. Similarly, classification results remain largely the same, except in the scenario with 45% contamination where the number of false-positives does increase.

## Acknowledgments

I thank Mateja Hajdinjak, Fernando Racimo, Cosimo Posth, Fernando Racimo, Janet Kelso, Elena Zavala, Janet Kelso Svante Pääbo and other members of the MPI genetics department for helpful discussion. This study was funded by the Max Planck Society and the European Research Council (grant agreement number 694707).

## Data Availability

No novel data was generated for this project. The unpublished data for *Chagyrskaya 8* is used ahead of publication with permission from the authors.

## Code Availability

Code to reproduce all results is available at https://www.github.com/benjaminpeter/admixfrog-sims and https://www.github.com/benjaminpeter/admixfrog

---

**Legend for Supplemental Table 1: All called introgressed fragments:** sample: Specimen fragment was called in. type: whether call is homozygous (‘homo’) or any ancestry (‘state’). Target: Source ancestry of fragments. Three-letter abbreviations are described in text. chrom: chromosome, pos, pos_end, pos_len: start, end position and length (in BP). Map, map_end, and map_len: start, end position and length (in cM); genes: Genes found in the introgressed fragment.

**Extended Data Table 1:** Data - Sample Summary Table. target: one of NEA: for Neandertal ancestry, NEA(h): Homozygous Neandertal ancestry; DEN: Denisovan ancestry. NEA*/DEN*: Ancestry using fixed prior. n(A/X): number of inferred autosomal/ X-chromosome fragments; Tot: Total amount of introgressed material in cM and Mb; Max: Length of longest tract; Gen, Years: Mean estimated age of admixture tracts in generations/years (rounded to 100).

| Specimen | Target | Age Range | n(A/X) | Tot(cM) | Tot(Mb) | Max(cM) | Max(Mb) | Gen | Years | Reference |
| --- | --- | --- | --- | --- | --- | --- | --- | --- | --- | --- |
| Denisova 2 | NEA | 122,700–194,400 | 95/1 | 212.6 | 173 | 25.72 | 17.6 | 50±10 | 1400±300 | 9 |
| Denisova 8 | NEA | 105,600–136,400 | 54/1 | 258 | 210 | 22.57 | 23.72 | 22±6 | 600±200 | 8 |
| Denisova 5 | DEN | 90,900–130,000 | 15/3 | 15.2 | 10.2 | 2.17 | 2 | 155±73 | 4500±2100 | 5 |
| Denisova 5 | DEN* | 90,900–130,000 | 18/3 | 14.1 | 10 | 2.17 | 2 | 212±92 | 6100±2700 | 5 |
| Denisova 3 | NEA* | 51,600–76,200 | 58/0 | 32.6 | 21.5 | 4.61 | 5.05 | 276±72 | 8000±2100 | 3 |
| Denisova 11 | NEA(h) | 115,700–140,900 | 51/0 | 16.6 | 13.3 | 1.01 | 1.12 | 795±223 | 23100±6500 | 6 |
| Denisova 11 | DEN(h) | 115,700–140,900 | 0/0 | 0 | 0 | - | - | - | - | 6 |
| Chagyrskaya 8 | DEN | 71,000–87,000 | 12/0 | 3.8 | 4.6 | 0.83 | 1.15 | 839±484 | 24300±14100 | 33 |
| Mezmaiskaya 1 | DEN | 60,000–70000 | 1/0 | 0.5 | 0.8 | 0.46 | 0.83 | - | - | 5 |
| Mezmaiskaya 2 | DEN | 42,960–44,600 | 2/0 | 0.5 | 1.1 | 0.24 | 0.83 | - | - | 29 |
| Goyet Q56-1 | DEN | 42,080–43,000 | 3/0 | 0.7 | 1.6 | 0.32 | 1.29 | - | - | 29 |
| Vindija 33.19 | DEN | 43,100–47,600 | 0/0 | 0 | 0 | - | - | - | - | 4 |
| HST | DEN | 62,000–183,000 | 0/0 | 0 | 0 | - | - | - | - | 20 |
| Les Cottés Z4-1514 | DEN | 42,720–43,740 | 2/0 | 0.4 | 1.2 | 0.25 | 1.17 | - | - | 29 |
| Spy 1 | DEN | 37,876–39,154 | 1/0 | 0.3 | 1.4 | 0.28 | 1.41 | - | - | 29 |
| Scladina | DEN | 95,000–173,000 | 0/0 | 0 | 0 | - | - | - | - | 20 |
| Denisova 4 | NEA | 55,200–84,100 | 0/0 | 0 | 0 | - | - | - | - | 8 |

**Extended Data Figure 1.**
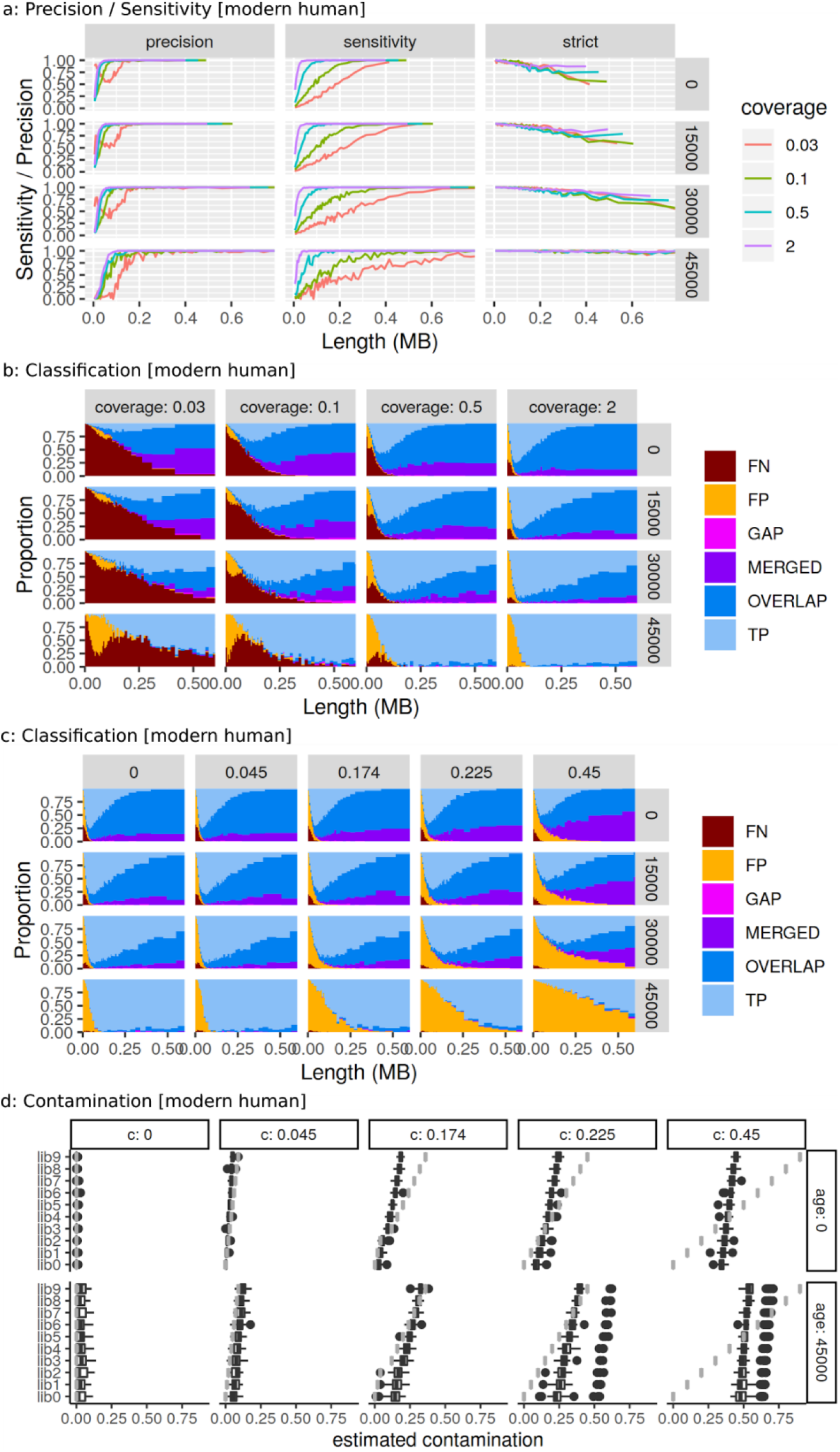
Simulations in human scenario for admixture 50,000 years ago. **a:** Precision, sensitivity and strict positive classification depending on coverage. **b:** Classification depending on sampling time (panel rows) and coverage (panel columns). TP: True Positives, FN: False Negatives, FP: False Positives. GAP: Single fragment inferred as multiple fragments. MERGED: Disjoint fragments inferred as a single fragment. OVERLAP: Multiple overlapping fragments inferred as single fragment (see Methods). **c:** Classification depending on sampling time (panel rows) and contamination rate (panel columns) for 2x samples. **d:** Contamination estimates for the ten simulated libraries for all five scenarios (panel columns). Grey lines give simulated contamination rates, boxplots give estimates for each library.

**Extended Data Figure 1.**
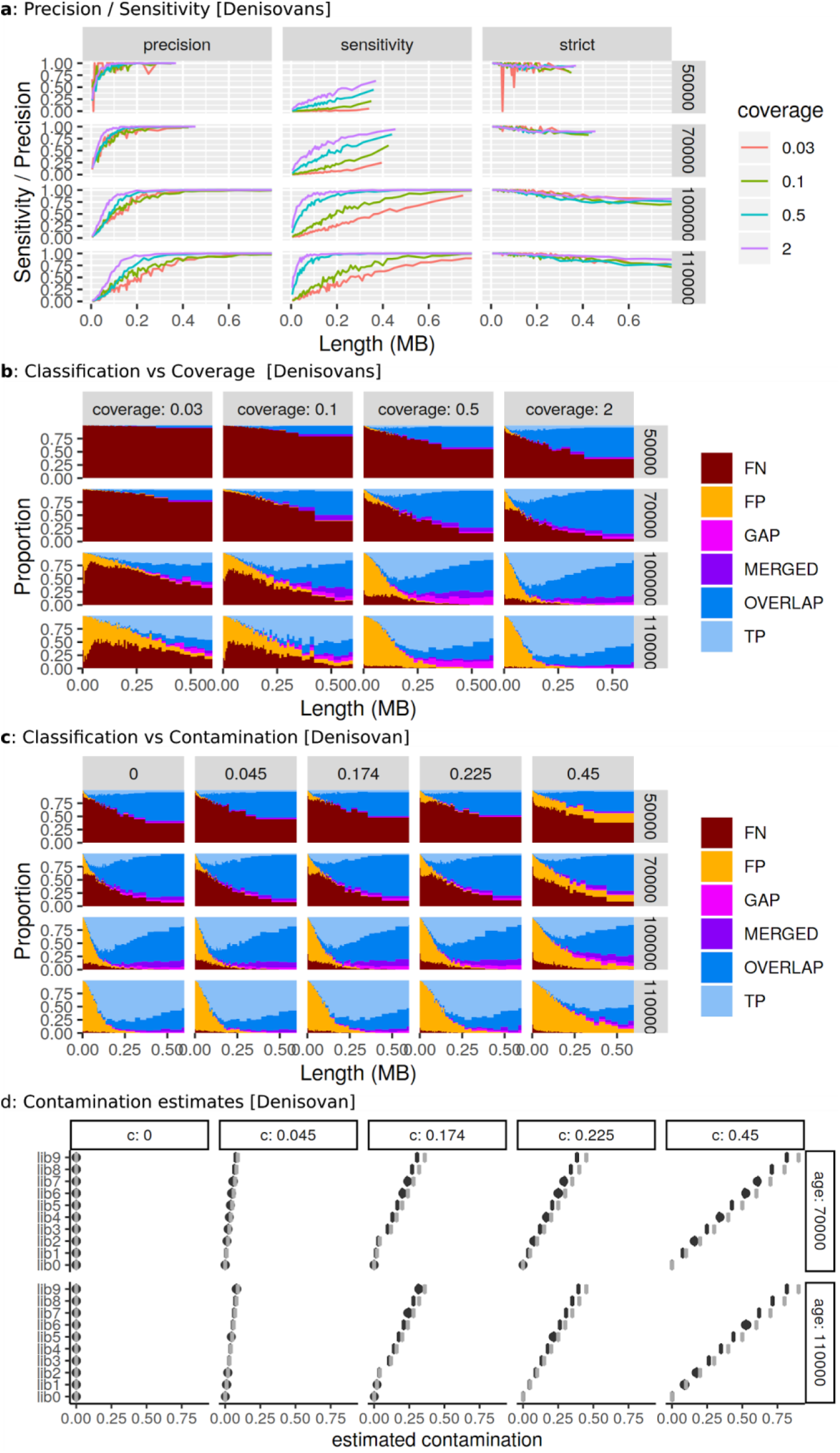
Simulations in archaic scenario for admixture 120,000 years ago. **a:** Precision, sensitivity and strict positive classification depending on coverage. **b:** Classification depending on sampling time (panel rows) and coverage (panel columns). TP: True Positives, FN: False Negatives, FP: False Positives. GAP: Single fragment inferred as multiple fragments. MERGED: Disjoint fragments inferred as a single fragment. OVERLAP: Multiple overlapping fragments inferred as single fragment (see Methods). **c:** Classification depending on sampling time (panel rows) and contamination rate (panel columns) for 2x samples. **d:** Contamination estimates for the ten simulated libraries for all five scenarios (panel columns). Grey lines give simulated contamination rates, boxplots give estimates for each library.

**Extended Data Figure 3.**
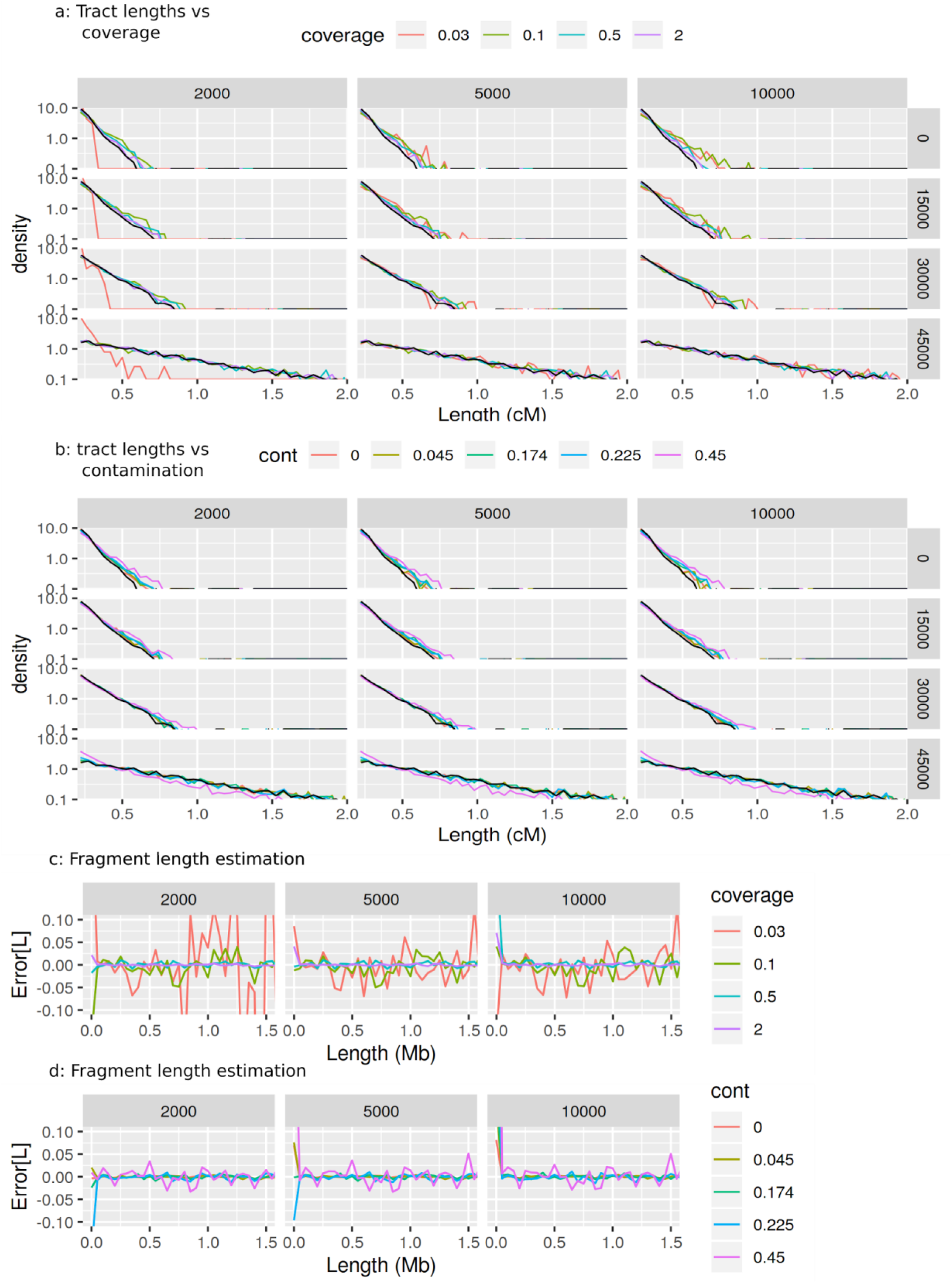
Simulations - Fragment length estimation in human scenario. **a:** Simulated (black) vs estimated distribution of inferred admixture fragment lengths depending on bin size (panel columns, in kb), age (panel rows, in years) and coverage (color). Densities are given on log-plot, so an exponential distribution appears as a line. The distribution is truncated at 0.2cM. **b:** Same as a, with contamination at 2x coverage. **c, d:** Mean error in fragment length estimation depending on bin size for a sample taken 30,000 years before present.

**Extended Data Figure 4:**
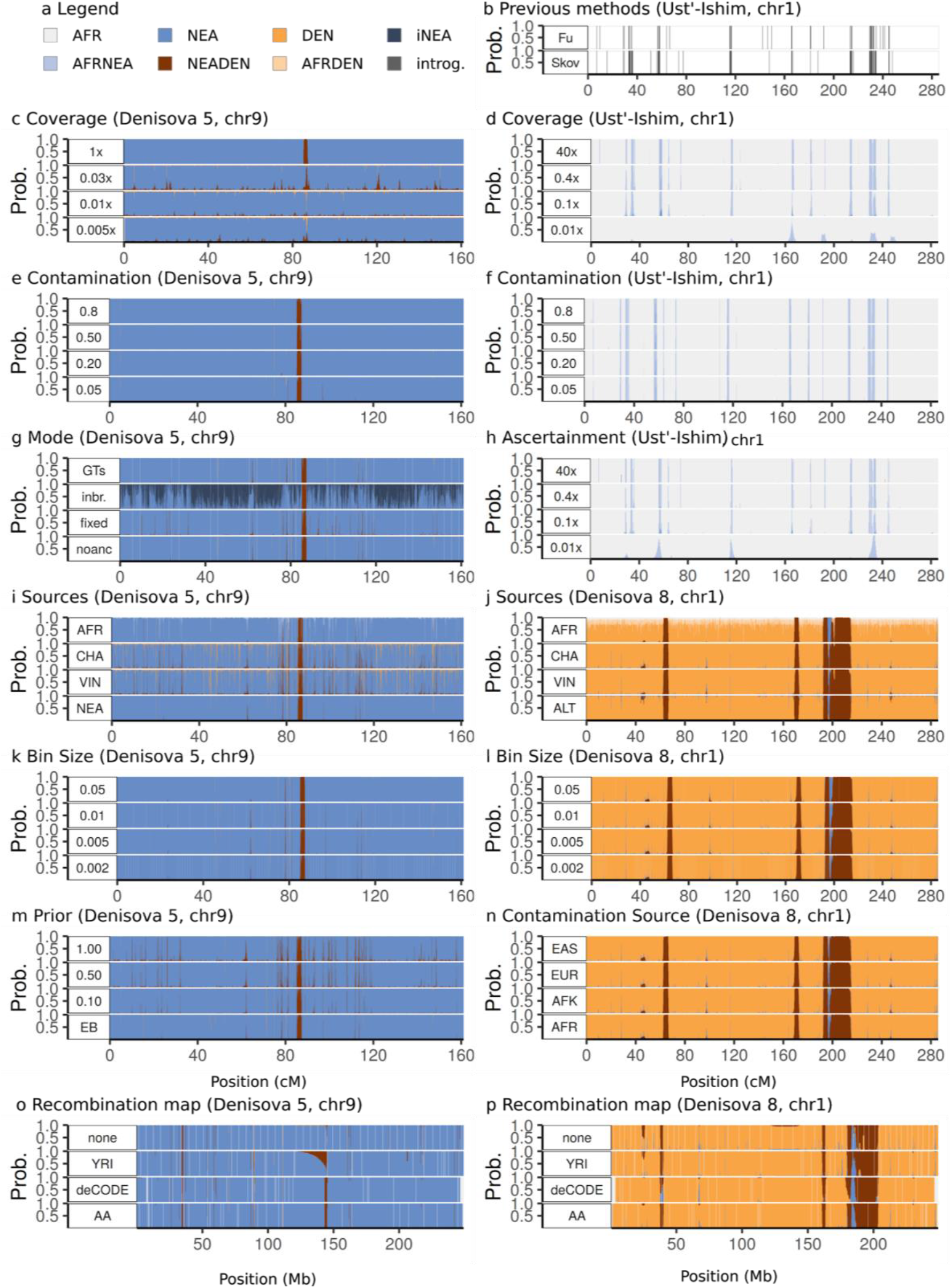
Empirical Experiments. I show posterior decodings of chr9 of *Denisova 5* (c, e, g, i, k, m, o) chr1 of *Ust’-Ishim* (b, d, f, h) and *Denisova 8* (j, l, n, p), and under varying conditions. **a:** Color legend; **b:** Inferences using previous approaches^30,48^; **c/d:** Downsampling to lower coverages **e/f:** adding contamination from the given contamination panel. **g**: inference using additional options: GTs: using called genotypes instead of read data; inbr: adding states for inbreed ancestry; fixed: no estimation of *F*/◻; noanc: no ancestral allele; error estimation; error: with error estimation; fix: All drift parameters fixed a priori. GTs: Inference done using called genotypes. **h**: SNP ascertainment: pARC: polymorphic in archaics (used for most analyses); 1240: modern human array from^49^; 3.7M: full array from ^49^; AAdm: archaic array from ^49^; **i/j**: Alternative sources using a single Neandertal (VIN/CHA/ALT), all Neandertals (NEA) or allowing AFR as an additional source (AFR). **k/l:** Size of bin (in cM). **m:** Fixing prior *a*_0_/*d*_0_ to 0.1, 0.5, 1 vs. estimating it from data (EB). **n:** Contamination panel EAS: East Asians, EUR Europeans AFR: Sub-Saharan Africans (from SGDP). AFK: Sub-Saharan Africans (from 1000g^54^). **o/p:** Effect of recombination map on inference: using no map (‘none’), HapMap-map using YRI^55^, deCODE-map^50^ and (AA) African-American map^43^.

**Extended Data Figure 5:**
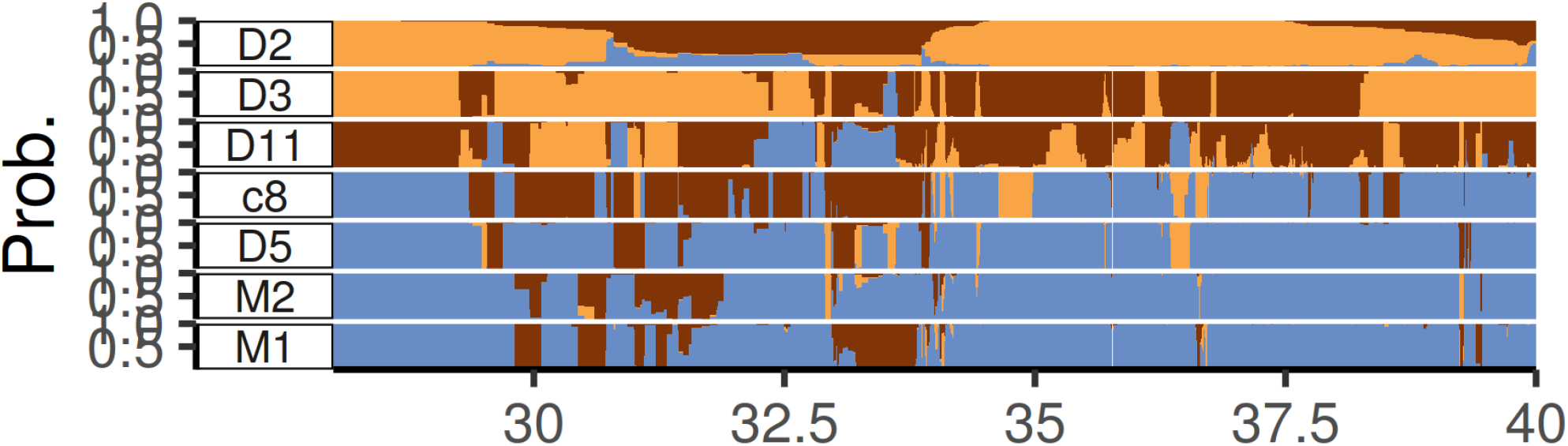
MHC locus on chromosome 6. I show the admixfrog decoding for chr6:28,000,000-40,000,000, a region overlapping the MHC region (chr6:28,477,797-33,448,354) and where an excess of admixture signals, in both directions, are detected. Given the prevalence of balancing selection and other potentially confounding issues, the admixture history of this locus can currently not be resolved.

**Extended Data Figure 6:**
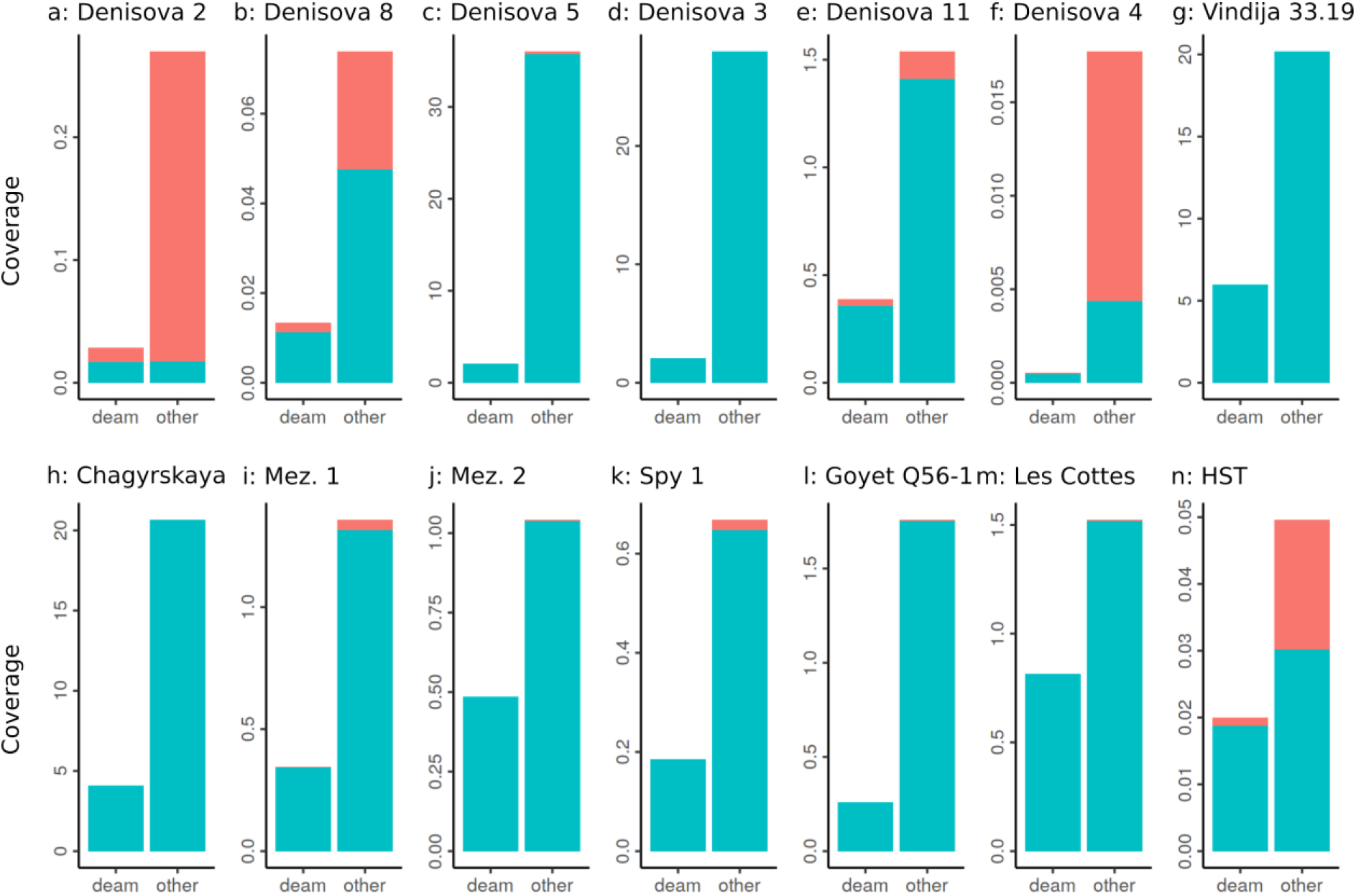
Data - Contamination estimates and coverage for samples analyzed in this study. Blue: estimated endogenous coverage. Red: Estimated contaminant coverage. Left bar represents reads carrying a deamination in the first three base pairs (‘deam’), right bar all other reads.

**Extended Data Figure 7:**
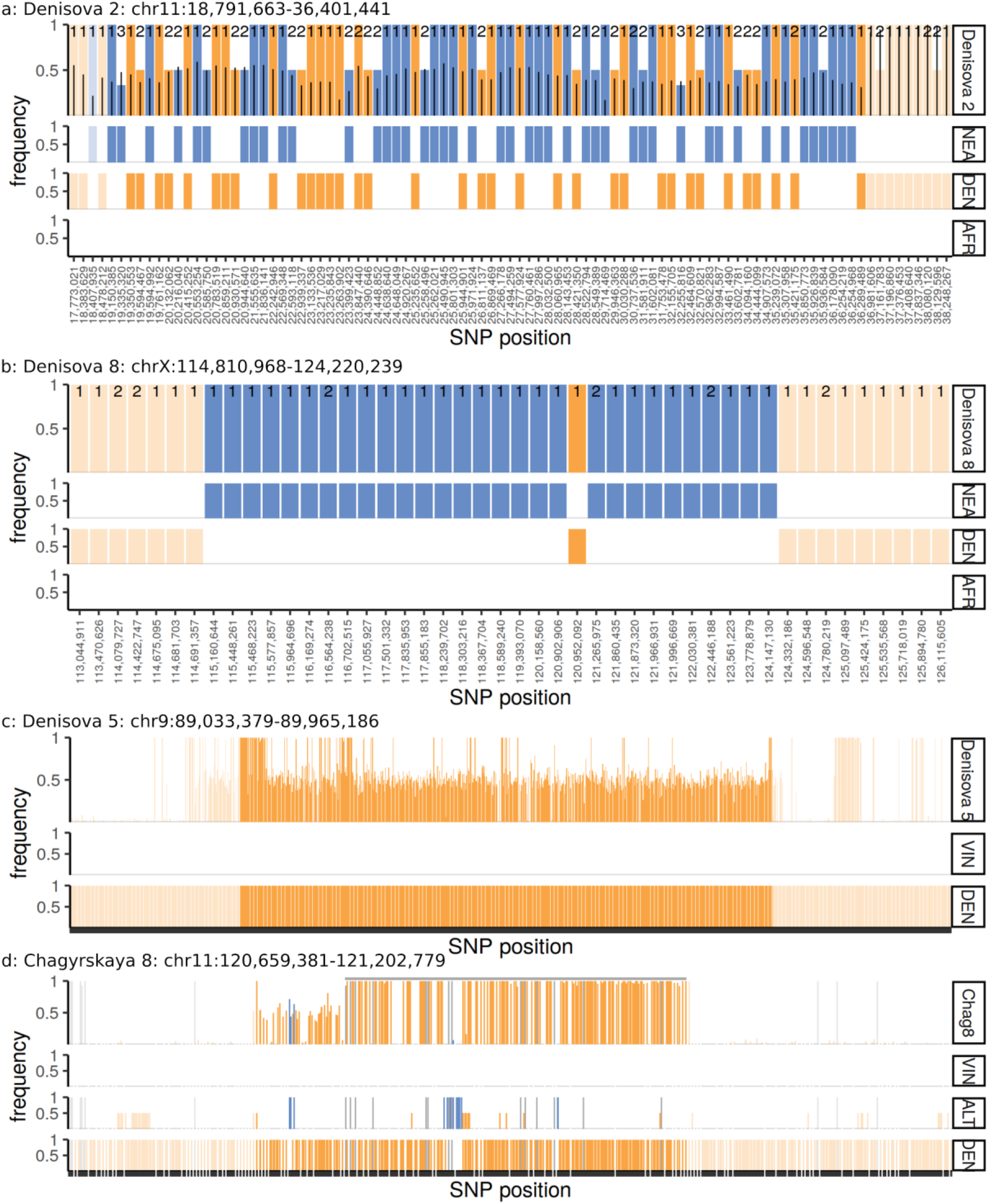
Validation of tracts. Shown are a subset of informative SNPs in inferred ancestry tracts. SNP fixed in Neandertals and Denisovans are colored blue and orange, respectively. The inferred tract is displayed in saturated colors; flanking regions are faded. In each panel, the top row displays the proportion of derived allele reads in the target genome. Numbers give the total number of reads for low-coverage genomes. Black line in *Denisova 2* indicates the posterior expected allele frequency in *Denisova 2*. Other rows give allele frequency in reference panels. **a *Denisova 2***: Only SNPs where at least one non-AFR read is present in *Denisova 2*, and where DEN and NEA are differentially fixed are plotted. **b***Denisova 8*: Same ascertainment as for *Densiova 2*. **c***Denisova 5*: Displayed are all SNP fixed between *Vindija 33.1*9 and *Denisova 3*, and the reads in *Denisova 5*. **d***Chagyrskaya 8*: A partially homozygous introgressed region on Chromosome 11, shown are all SNPs where Altai or *Denisova 3* are fixed for an allele that differs from Vindija 33.19.

**Extended Data Figure 8:**
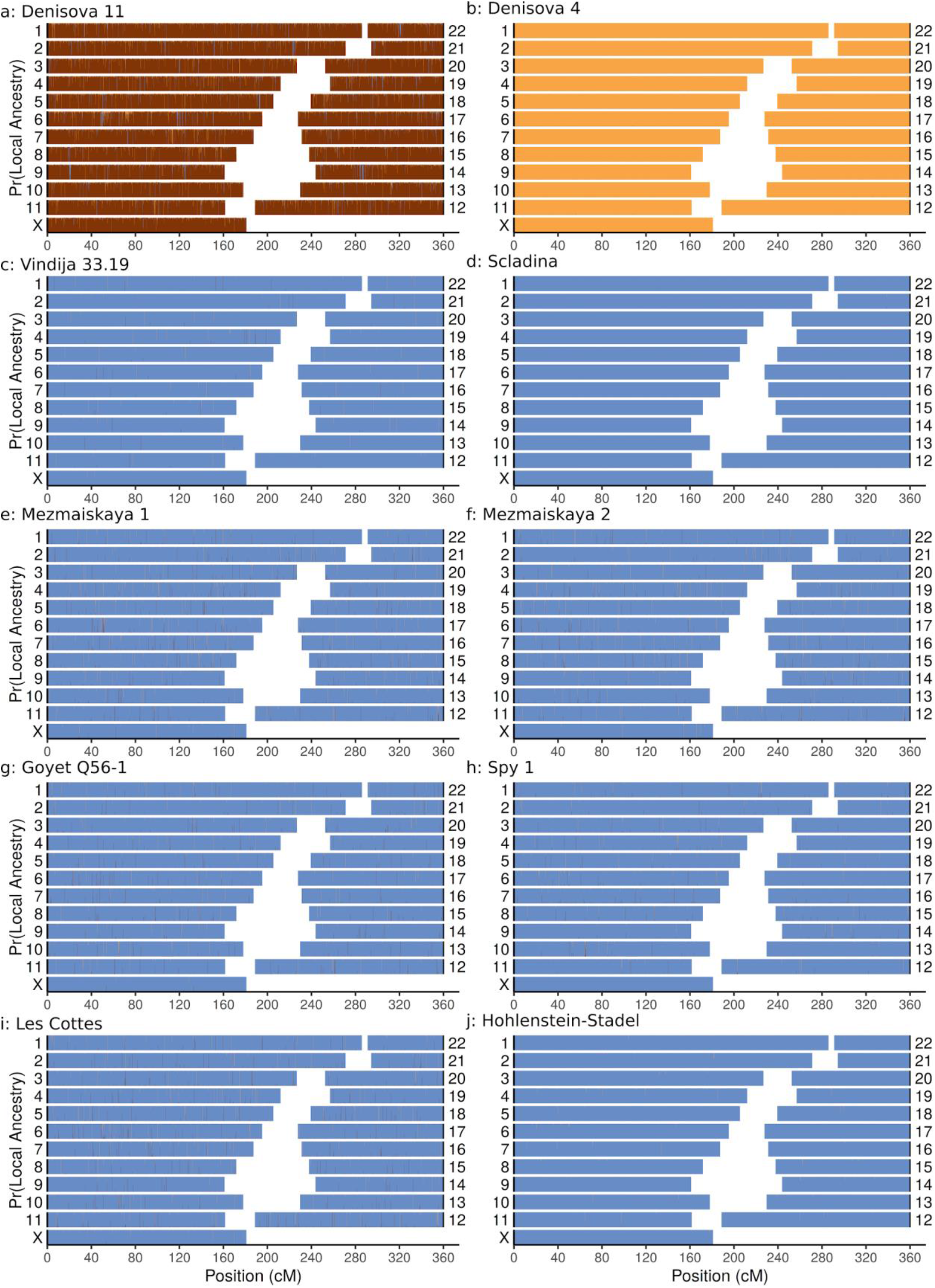
Admixfrog local ancestry posterior decoding for other archaic genomes. Homozygous Denisovan ancestry, homozygous Neandertal ancestry and heterozygous ancestry are in orange, blue and brown, respectively.

**Extended Data Figure 9:**
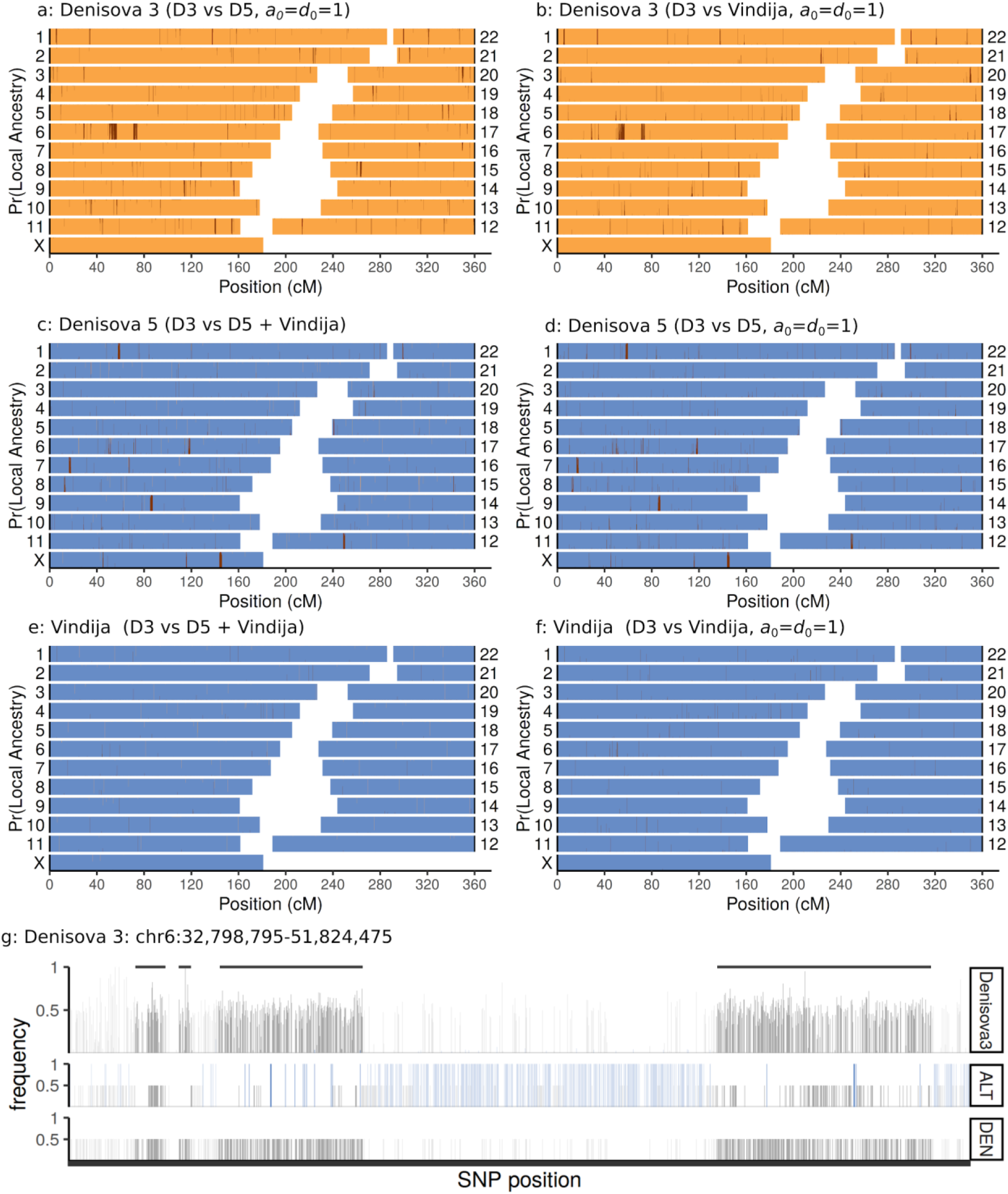
*Denisova 3* overview. Admixfrog posterior decodings of *Denisova 3* (a,b), *Denisova 5* (c, d) and Vindija 33.19 (e, f) using a fixed (a, b, d, f) and empirical Bayes prior (c, e). Homozygous Denisovan ancestry, homozygous Neandertal ancestry and heterozygous ancestry are in orange, blue and brown, respectively. For the Neandertals, results are consistent between analyses, but more noisy using the fixed prior. **g:** Four fragments on chr6 that are candidates for introgression, due to the high number of heterozygous sites and absence of fixed differences between Neandertals and *Denisova 3*. Called fragments are marked with grey horizontal lines.

