## Supplemental Text 1 for "100,000 years of gene flow between Neandertals and Denisovans in the Altai mountains"

### Supplement 1: Admixfrog Model Details

Benjamin Peter

February 8, 2020

#### Model Overview

Here, we present the model and algorithms behind `admixfrog` in full detail. The Hidden Markov Model (HMM) implemented in `admixfrog` classifies a single *target* genome into regions that were derived from different diverged "source" populations. This is motivated by the admixture between Denisovans, Neandertals and modern humans, although the model is applicable to many other taxa.

We subdivide the diploid target genome into small bins of fixed length (either in genetic map units or physical size ). For each bin, we define a latent variable $Z_l$ that designates the combination of ancestry sources. For example, in the above scenario, we would have three homozygous ancestries (modern human, Neandertal and Denisovan), as well as three heterozygous ancestries (all possible pairs of above ancestries). The vector of all $Z_l$ will be denoted as $\mathbf{Z}$, and the space of all vectors as $\mathcal{Z}$.

In each bin, there may be zero or more biallelic observed SNPs, whose genotype is denoted by $G_{ls} \in \{0, 1, 2\}$. We assume that SNPs in different bins are independent conditional on $\mathbf{Z}$. Within each bin, we similarly assume that SNPs are independent conditional on $Z_l$. For a particular SNP, $P(G_{ls}|Z_l)$ will depend on the allele frequency $p_{sk}$ in the relevant source $k$, which is estimated from a reference sample from the source. The vector of all genotypes is denoted as $\mathbf{G}$, and the space of all such vectors is $\mathcal{G}$.

Finally, we assume that for each SNP we have zero or more reads sampled from $R$ disjoint read groups. The total number of reads from a particular read group at a particular SNP $G_{sl}$ is $n_{rsl}$, and the random variable denoting the number of non-reference alleles is $O_{rsl}$. Each read group will have its own contamination rate $c_r$ and error parameter $e_r$, that are estimated directly from the data.

We are primarily interested in estimating the latent states $\mathbf{Z}$ and $\mathbf{G}$, but we also estimate the transition matrix $A$ between states (which, in turn, is informative about admixture proportion and times), the contamination and error rate for each read group, the substructure in each source $\tau_k$, and the average drift since admixture from each source $F_k$.

#### Notation overview

To summarize, the notation is as follows:

- $R, L$ denote the number of read groups, and hidden-state bins, respectively
- $K$ is the number of sources
- $S_l$ is the number of SNPs in bin $l$
- $n_{rsl}$ the number of reads of readgroup $r$ at SNP $s$ of bin $l$.
- $\mathbf{O} = (O_{rsl})$ the set of derived-alleles from library $r$ at SNP $s$ in bin $l$
- $\mathbf{G} = (G_{sl})$ the set of genotypes of SNP $s$ in bin $l$
- $\mathbf{Z} = (Z_l)$ be the sequence of hidden states

- $a_{sli}, d_{sli}$ the number of ancestral and derived allele counts in reference $i$ at $\text{SNP}_{sl}$, (or more generally parameters of a Beta prior)
- $A$ be a transition matrix
- $F_k$ a parameter estimating coalescence sind gene flow from source $k$.
- $\tau_k$ measuring genetic drift between gene flow and the reference sample from source $k$.
- $c_r, e_r$ proportion of contaminant reads and error rate in read group $r$
- $p_{slk}$ the allele frequency of source $k$ at position $sl$.
- $p^c_{sl}$ the allele frequency of a contaminant at position $sl$.
- $\theta = (AF_k, \tau_k, e_r, c_r)$, the set of all parameters to be estimated

The model can be summarized as

- $O_{rsl}|G_{sl}, n_{rsl} \sim \text{Binomial}(O_{rsl}; n_{rsl}, p(e_r, c_r, p^c_{sl}, G_{sl}))$
- $\psi_k \sim \text{Bernoulli}(F_k)$
- $p_{slk}|Z_l = k \sim \text{Beta}(a_k\tau_k, d_k\tau_k)$
- $G_{sl}|Z_l = k \sim \psi_k\text{Binomial}(1, p_{slk}) + (1-\psi)\text{Binomial}(2, p_{slk})$
- $Z_s|Z_{s-1} = k' \sim \text{Categorical}(A_{k'})$

### Algorithm details

We assume basic familiarity in the techniques and derivation of the standard algorithms for HMMs. Readers unfamiliar might consider standard text books, e.g (Durbin et al., 1998). The complete-data likelihood (assuming the latent variables $\mathbf{G}, \mathbf{Z}$ are known) is maximized using an EM-algorithm. The complete data log-likelihood is

$$\begin{aligned}\mathcal{L} = \log P(\mathbf{O}, \mathbf{Z}, \mathbf{G}|\theta) &= \sum_{r=1}^{R}\sum_{s=1}^{S_l}\sum_{l=1}^{L} \log P(O_{rsl}|G_{sl}, c_r, e_r) \\ &+ \sum_{l=1}^{L}\sum_{s=1}^{S_l} \log P(G_{sl}|Z_l = k, F_{Z_l}, \tau_{Z_l}) \\ &+ \sum_{l=1}^{L} \log P(Z_l|Z_{l-1}, A) + \log P(Z_0). \end{aligned} \tag{1}$$

#### Transition and initial probabilities

The transition probabilities $P(Z_l|Z_{l-1}, A)$ follow a homogeneous Markov chain based on the transition matrix $A$.

#### State Likelihood

In contrast to standard HMMs, our emissions involve the unknown genotypes as an additonal latent variable. We refer to $P(G_{sl}|Z_l = k, F_{Z_l}, \tau_{Z_l})$ as the state likelihood, and $P(O_{rsl}|G_{sl}, c_r, e_r)$ as the genotype likelihood. The state likelihood $P(G_{sl}|Z_l = k, a_k, d_k, F_k, \tau_k)$ represents the probability of the genotype at a single locus, given the hidden state $Z_l$ (i.e ancestry). The state likelihood depends on the allele frequencies in the samples from the putative sources, and we take into account three sources of uncertainty:

1. Sampling uncertainty in the reference panel

2. Genetic drift in the target population after introgression
3. Population substructure and drift in the source populations

The resulting probabilities differs slightly between homozygous and heterozygous ancestries, and so we derive them sepearately.

**Homozygous Hidden States**

As we deal with a single hidden-state bin, locus, and source, subscripts designating them are omitted in this section. We start with the simplest possible model, and add complexity in sequence.

**Known allele frequency** If, the allele frequency $p$ is known,

$$G \sim Binom(2, p).$$

**Sampling uncertainty** As $p$ is unknown, we estimate it from a sample where we observe $a'$ ancestral and $d'$ derived alleles, respectively. In this case we model the allele frequency as

$$p \sim Beta(d' + d_0, a' + a_0)$$

The prior $d_0, a_0$ can be justified by noting that even if we only observe ancestral alleles in our sample, there is still a small chance that some individuals might have a derived allele, particularly when the sample size is small. There are differing opinions about the values of $d_0, a_0$. For large samples, Balding and Nichols (1995) proposed a uniform prior, i.e. $a_0 = d_0 = 1$. By noting that the derived allele occurs proportionally to $\theta/p$, one can justify $d_0 = 0, a_0 = 1$ (Racimo et al., 2016). The third option we use here is an empirical Bayes prior (Efron, 2012): In the absence of data at a particular locus, the genome wide site-frequency spectrum is a sensible guess for the allele frequency distribution of the focal locus. Hence, we obtain an empirical prior by modeling the (folded or unfolded) site-frequency spectrum as a $Beta(d_0, a_0)$ distribution. $d_0$ and $a_0$ are fitted using a method-of-moments approach from the genome-wide distribution of derived allele frequencies.

Writing $d = d' + d_0, a = a' + a_0$, the probability of $G$ is then

$$\begin{aligned} G &\sim Betabinom(2, d, a) \\ P(G|d, a) &= \binom{2}{G} \frac{B[G + d, 2 - G + a]}{B[d + a]} \end{aligned} \quad (2)$$

where $B[.]$ denotes the beta function.

**Genetic drift in target** In many cases, the target individual will not be sampled immediately after admixture, but thousands of generations after. Thus, we need to take into account that the allele frequency might have changed.

Conceptually, genetic drift will increase the chance that the two alleles in the target are derived from the same ancestor at introgression time: If the target is very distant from the source, the probability $F$ that the two alleles are identical by descent approaches one. Conversely, if we have a very close reference, that probability is zero. In other words, $F$ is the probability of coalescence before gene flow. Therefore,

$$\begin{aligned} P(G = 2|a, d, F) &= \int P(G|p)P(p|a, d)df \\ &= \int \left[Fp + (1 - F)p^2\right] P(p|a, d)df \\ &= F\mathbb{E}\left[p|a, d\right] + (1 - F)\mathbb{E}\left[p^2|a, d\right] \end{aligned} \quad (3)$$

For the Balding-Nichols-model, we thus obtain

$$P(G = 2|a, d, F) = F\frac{d}{a + d} + (1 - F)\frac{d(d + 1)}{(a + d)(a + d + 1)}$$
$$= \frac{d^2 + d + adF}{(a + d)(a + d + 1)} \tag{4a}$$

We note that the allele frequency distribution only enters through the first two moments, and hence any model that would us give the some moments for the allele frequency in the source will give the same results. Furthermore, if $F > 0$, this will result in an excess of homozygous emissions, as expected by the loss of (ancestral) heterozygosity expected under genetic drift.

The probability for $G = 0$ is obtained by symmetry; we simply switching $a$ and $d$. The probability of a heterozygous genotype ($G = 1$) is obtained by subtracting them from one:

$$P(G = 1|a, d) = 2(1 - F)\Big[\mathbb{E}\left[p|a, d\right] - \mathbb{E}\left[p^2|a, d\right]\Big] = \frac{2ad(1 - F)}{(a + d)(a + d + 1)} \tag{4b}$$

**Drift and population structure in source** In many cases, the available samples from the source might be only distantly related to the true introgressing individual. Alternatively, ascertainment bias or non-random sampling might result in a lower uncertainty in the reference allele frequency than might be expected from the sample size. For example, we want to infer Neandertal introgression by using the Altai Neandertal (**?**) as a reference, despite it being substantially diverged from the introgressing Neandertal. The approach chosen here is to scaling the variance (but not mean) of the allele frequency distribution: For this purpose, we introduce a parameter $\tau$

$$p|a, d, \tau \sim Beta(d\tau, a\tau)$$

which leads to (starting from eq 3):

$$P(G = 2|a, d, F, \tau) = F\mathbb{E}[p|a, d, \tau] + (1 - F)\mathbb{E}[p^2|a, d, \tau]$$
$$= F\frac{d}{a + d} + (1 - F)\frac{d(d\tau + 1)}{(a + d)(a\tau + d\tau + 1)}$$
$$= \frac{\tau d^2 + d + adF\tau}{(a + d)(\tau a + \tau d + 1)} \tag{5a}$$
$$P(G = 1|a, d, F, \tau) = \frac{2ad(1 - F)\tau}{(a + d)(\tau a + \tau d + 1)} \tag{5b}$$

##### Heterozygous States

For a heterozygous latent state, the model is considerably simpler: We know that there is exactly one haploid genome sampled from each source. If we assume that the sources have allele frequencies $p_i$ (described by a Beta distribution with parameters $d_i, a_i$, respectively):

$$P(G_i = 1|\tau_i, F_i, d_i, a_i) = \int pP(f|d_i, a_i, p\tau_i, F_i)dp$$
$$= E[p|d_i, a_i, F_i, \tau_i]$$
$$= \frac{d_i}{a_i + d_i} \tag{6}$$

and hence is independent from both $F$ and $\tau$. Intuitively, this is because if we just sample one allele from a population, that population's allele frequency is the best guess for the state of our sample. Thus for sources with samples $(a_i, d_i)$ and $(a_j, d_j)$,

$$P(G=0) = \frac{a_i a_j}{(d_i + a_i)(d_j + a_j)} \tag{7a}$$

$$P(G=1) = \frac{a_i d_j + a_j d_i}{(d_i + a_i)(d_j + a_j)} \tag{7b}$$

$$P(G=2) = \frac{d_i d_j}{(d_i + a_i)(d_j + a_j)} \tag{7c}$$

##### Haploid States

On the sex chromosome, or when inbreeding is present, we further encounter haploid regions. Here, $G$ takes only values of 0 or 1, and reasoning analogous to the heterozygous case yields

$$P(G=0) = \frac{a}{a+d} \tag{8a}$$

$$P(G=1) = \frac{d}{a+d} \tag{8b}$$

#### Genotype Likelihood

We assume a simple Bernoulli mixture of contaminant and endogenous reads similar to e.g. Racimo et al. (2016). The error rate $e$ switches the allele to the other state.

$$P(O_r|G, c_r, n_r, e_r, p^c) \sim Binom(O_r; n_r, p) \tag{9}$$

where $p = (1-e_r)p' + e_r(1-p')$ and $p' = c_r p_c + (1-c_r)G$.

### Parameter estimation

One of the strengths of this model formulation is that all parameters can be efficiently estimated from the data. Here, we lay out the specific M-steps implemented in `admixfrog`.

#### Estimationg Transitions and Initial State

As the model is a homogeneous Hidden-Markov model, $A$ can be estimated using the standard Baum-Welch algorithm. The initial distribution $P(Z_0) = \alpha_0$ is set to the stationary distribution of $A$ after each iteration. The hierarchical nature of eq. 1 simplifies optimization considerably. To estimate $\tau$ and $F$, only the terms $\log P(G|Z)$ are needed, as all other terms are independent of $\tau$ and $F$. Likewise, to estimate $c$ and $e$, only $\log P(O|G)$ is required.

#### Estimating F and $\tau$

The $Q$-function (dropping the terms not depending on $F$ and $\tau$)is

$$\begin{aligned} Q(F,\tau|F',\tau') &= E[\log P(\mathbf{O},\mathbf{Z},\mathbf{G}|F,\tau)P(\mathbf{G},\mathbf{Z}|\tau',F',\mathbf{O})] \\ &= \sum_{\mathbf{Z}\in\mathcal{Z},\mathbf{G}\in\mathcal{G}} \log P(\mathbf{G}|\mathbf{Z},F,\tau)P(\mathbf{G}|\mathbf{Z},\mathbf{O},F',\theta')P(\mathbf{Z}|\mathbf{O},F',\tau') \\ &= \sum_k \sum_{g=0}^{2} \sum_{l=1}^{L} \sum_{s=1}^{S_l} \log P(G_{sl}=g|Z_l=k,F_k,\tau_k)P(G_{sl}=g|Z_l=k,F',\tau',\mathbf{O})P(Z_l=k|\mathbf{O},F',\tau') \end{aligned} \tag{10}$$

Since both $F_k$ and $\tau_k$ only depend on the terms for the respective homozygous hidden states, we numerically optimize

$$(\hat{F}_k,\hat{\tau}_k) = \underset{F,\tau}{\operatorname{argmax}} \left[\sum_{g=0}^{2} \sum_{l=1}^{L} \sum_{s=1}^{S_l} \log P(G_{sl}=g|Z_l=k,F,\tau)P(G_{sl}=g|Z_l=k,F',\tau',\mathbf{O})P(Z_l=k|\mathbf{O},F',\tau')\right] \tag{11}$$

where $P(Z_l = k|\mathbf{O}, \theta')$ is the output of the forward-backward algorithm, $P(G_s|Z_l = k, \tau_k, F_k)$ is given by eq. 5, and

$$P(G_{sl}|Z_l, \mathbf{O}, F', \tau') = \frac{P(O_{sl}|G_{sl}, c')P(G_{sl}|Z_l, F', \tau')}{\sum_{g=0}^{2} P(O_{sl}|G_{sl} = g, c')P(G_{sl} = g|Z_l, F', \tau')} \tag{12}$$

This equation follows by applying Bayes theorem:

$$\begin{aligned} P(G, Z|\mathbf{O}) &= P(G|Z, \mathbf{O})P(Z|\mathbf{O}) \\ &= P(G|Z, O_l)P(Z|\mathbf{O}) \\ &= \frac{P(O_l|G)P(G|Z)}{P(O_l|Z)} P(Z|\mathbf{O}) \end{aligned} \tag{13}$$

as $G_l|Z_l$ is independent of all observations except $O_l$.

#### Estimating Contamination and Error

The update for contamination and error per read group are analogous:

$$\begin{aligned} Q(c, e|c', e') &= E[\log P(\mathbf{O}, \mathbf{Z}, \mathbf{G}|c, e)P(\mathbf{G}|\mathbf{Z}, \theta')P(\mathbf{Z}|\theta')] \\ &= \sum_{\mathbf{Z} \in \mathcal{Z}} \sum_{\mathbf{G} \in \mathcal{G}} \log P(\mathbf{O}|\mathbf{G}, c, e)P(\mathbf{G}|\mathbf{Z}, \theta')P(\mathbf{Z}|\mathbf{O}, \theta') \\ &= \sum_{r=1}^{R} \sum_{k} \sum_{g=0}^{2} \sum_{s=1}^{S_l} \sum_{l=1}^{L} \log P(O_{rsl}|G_{sl} = g, c_r, e_r)P(G_{sl}|Z_l = k, \theta', O)P(Z_l = k|\mathbf{O}, \theta') \end{aligned} \tag{14}$$

We thus optimize for each read group

$$(\hat{c}_r, \hat{e}_r) = \operatorname*{argmax}_{e_r, c_r} \sum_{k} \sum_{g=0}^{2} \sum_{s=1}^{S_l} \sum_{l=1}^{L} \log P(O_{rsl}|G_{sl} = g, c_r)P(G_{sl} = g|Z_l = k, \theta', \mathbf{O})P(Z_l = k|\mathbf{O}, \theta') \tag{15}$$

using (9) and (12).

Richard Durbin, Sean R. Eddy, Anders Krogh, and Graeme Mitchison. *Biological sequence analysis: probabilistic models of proteins and nucleic acids*. Cambridge university press, 1998.

Bradley Efron. *Large-Scale Inference: Empirical Bayes Methods for Estimation, Testing, and Prediction*. Cambridge University Press, November 2012. ISBN 978-1-139-49213-3. Google-Books-ID: kfE7VkqbGawC.

Fernando Racimo, Gabriel Renaud, and Montgomery Slatkin. Joint Estimation of Contamination, Error and Demography for Nuclear DNA from Ancient Humans. *PLOS Genetics*, 12(4): e1005972, April 2016. ISSN 1553-7404.
